# LegNet: a best-in-class deep learning model for short DNA regulatory regions

**DOI:** 10.1101/2022.12.22.521582

**Authors:** Dmitry Penzar, Daria Nogina, Elizaveta Noskova, Arsenii Zinkevich, Georgy Meshcheryakov, Andrey Lando, Abdul Muntakim Rafi, Carl de Boer, Ivan V. Kulakovskiy

## Abstract

**Motivation:** The increasing volume of data from high-throughput experiments including parallel reporter assays facilitates the development of complex deep learning approaches for DNA regulatory grammar.

**Results:** Here we introduce LegNet, an EfficientNetV2-inspired convolutional network for modeling short gene regulatory regions. By approaching the sequence-to-expression regression problem as a soft classification task, LegNet secured first place for the autosome.org team in the DREAM 2022 challenge of predicting gene expression from gigantic parallel reporter assays. Using published data, here we demonstrate that LegNet outperforms existing models and accurately predicts gene expression *per se* as well as the effects of single-nucleotide variants. Furthermore, we show how LegNet can be used in a diffusion network manner for the rational design of promoter sequences yielding the desired expression level.

**Availability and Implementation:** https://github.com/autosome-ru/LegNet. The GitHub repository includes the Python code under the MIT license to reproduce the results presented in the study and a Jupyter Notebook tutorial.

**Supplementary Information:** Online-only supplementary data are available at Bioinformatics online.

## Introduction

The basic level of gene expression regulation in eukaryotes, the mRNA transcription, is controlled by transcription factors (TFs), which bind cis-regulatory regions, promoters, and enhancers, and affect the assembly and functioning of the mRNA transcription machinery (Wasserman and Sandelin, 2004). The transcription factors can recognize specific DNA patterns, allowing them to act at various genomic addresses and affect particular sets of target genes (Lovering *et al*., 2021). It is a longstanding challenge in computational biology to decipher the sequence-level regulatory code, from predicting individual TF binding sites of varying affinity to identifying composite elements (Kel-Margoulis *et al*., 2002) and completing sequence-level annotation of promoters and enhancers.

A commonly accepted approach is bottom-up, where binding specificities of individual transcription factors are profiled with various TF-centric techniques (Jolma *et al*., 2013), revealing TF-specific binding motifs. With individual motifs at hand, the higher-order regulatory grammar can also be studied *in silico* (Boeva, 2016). However, genome-level analysis is hampered by numerous confounding factors, hence multiple direct experiments are required to explicitly profile the binding preferences of TF complexes (Isakova *et al*., 2017; Jolma *et al*., 2010). It remains challenging to apply the knowledge obtained in vitro to genomic regulatory regions and performing direct experiments for all TF combinations also remains hardly realistic.

A possible alternative approach to resolving the rules of regulatory grammar comes with massive parallel reporter assays (Klein *et al*., 2020), which can profile the activity of dozens of millions of synthetic or genomic regulatory sequences in a single experiment (de Boer *et al*., 2020; Sahu *et al*., 2022). The resulting data are uniform and diverse enough to allow an orthogonal approach: properly trained biochemical (de Boer *et al*., 2020) and advanced machine learning models (Vaishnav *et al*., 2022; Almeida *et al*., 2021) can provide quantitative and highly accurate predictions of regulatory activity just from the DNA sequence. In terms of machine learning, two questions remain unanswered in this setting. First, given the currently available data, if there remains enough space for further significant improvement of the computational models. Second, whether the high-level deep learning architectures such as attention transformers are truly necessary for modeling regulatory regions, or if the task can be handled by properly designed fully convolutional networks.

Here we introduce the LegNet convolutional network that our autosome.org team used to secure 1st place in all subchallenges of the DREAM 2022 challenge focused on predicting expression yield from a gigantic parallel reporter assay (GPRA) performed with yeast cells (**Figure 1**). Using previously published GPRA data, we demonstrate that LegNet outperforms existing methods in predicting both expression and sequence variant effects and highlight particular features of the model architecture that affect performance the most. Further, we demonstrate how LegNet can be used in diffusion generative modeling as a step toward the rational design of gene regulatory sequences.

**Figure 1.**
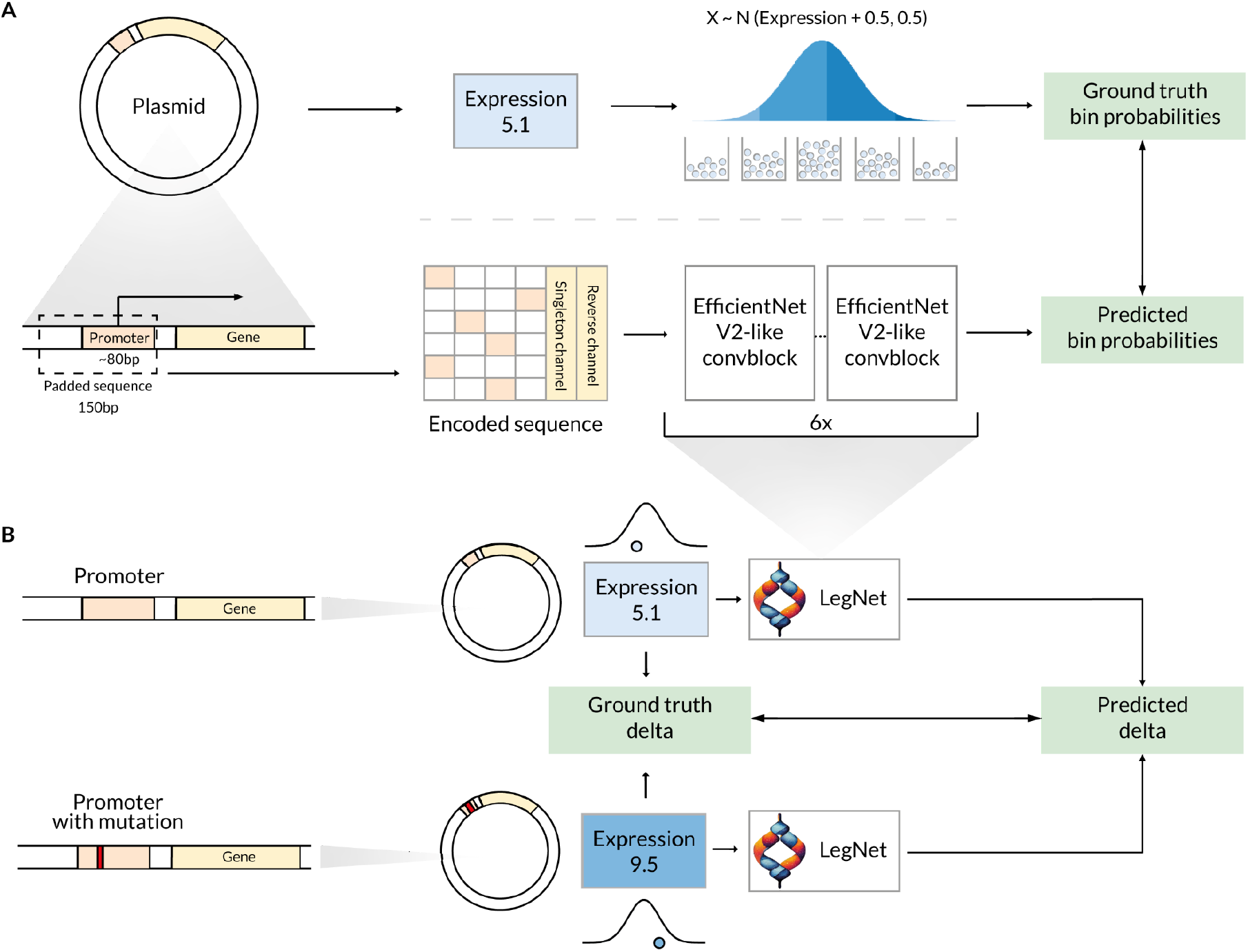
Learning and predicting promoter expression and effects of single-nucleotide variants from massive parallel reporter assays with LegNet. **A**. An overall pipeline. The regression task is reformulated as the soft-classification problem mirroring the original experimental setup where cells were sorted into different bins depending on reporter protein fluorescence. Bottom: sequence encoding and prediction of the expression bin probabilities with LegNet. **B**. Variant effect estimation with LegNet. Both original and mutated promoter sequences are passed separately to the trained neural network. The variant effect is estimated as a difference between corresponding predictions and compared against the ground truth experimental data.

## Methods

### Key ideas behind LegNet

Trending deep learning applications in biology are biased toward large-scale attention transformer models (Fudenberg *et al*., 2020; Avsec *et al*., 2021; Lin *et al*., 2022). Transformers indeed perform well in protein structure prediction and text mining, and particular applications of such models are successful in genomic data analysis (Fudenberg *et al*., 2020; Avsec *et al*., 2021). Yet, for the latter, it remains unclear whether it was the attention mechanism that contributed the most to their performance. Furthermore, depending on the test scenario, the quality of such models can be overestimated (Sasse *et al*., 2023), and they may in fact fail to capture long-range interactions (Karollus *et al*., 2022). In the context of parallel reporter assays with relatively short tested sequences, the application of a transformer-based approach might be an overcomplication compared to advanced fully convolutional networks. With this in mind, we took advantage of the EfficientNetV2 (Tan and Le, 2021) with its recent success in image analysis, and introduced several modifications to its architecture to account for specifics of the GPRA data.

### Experimental data overview

#### Yeast GPRA data of the DREAM2022 challenge

The initial version of LegNet was constructed to solve the 'sequence-to-expression' problem of the DREAM2022 challenge (Rafi *et al*., 2023) i.e. predicting promoter activity from GPRAs (Vaishnav *et al*., 2022). The challenge data consisted of nearly 6.7 million promoter-driven expression measurements in the yeast *S. cerevisiae* cultured in chardonnay grape juice. In the GPRA experiment, yeast cells were transformed with a plasmid containing the YFP gene controlled by 80bp random DNA sequences put into a promoter-like context, and the constitutively expressed RFP gene. Based on logarithmic relative protein fluorescence, yeast cells were sorted into 18 expression bins (numbered 0 to 17). The expression estimate for a particular promoter sequence is calculated as a weighted average of the numbers of expression bins where it was found (de Boer *et al*., 2020).

In the DREAM challenge, the train set consisted of 6.7 million random promoter sequences. The labels for the independent test set of 71 thousand promoters were hidden during the challenge and included the higher quality measurements obtained for various classes of promoters, including synthetic sequences and native yeast promoters. The details of the challenge setup, including the train and test data and the description of individual subchallenges, are described in detail in (Rafi *et al*., 2023).

For LegNet, we used the DREAM2022 challenge data (1) to build and verify the original model architecture, (2) to perform an ablation study for identifying key model elements contributing to the final performance the most, and (3) to analyze the performance of the LegNet-ensemble model.

#### Previously published yeast GPRA data

To further showcase LegNet applicability and verify its predictive performance, we used previously published GPRA results (Vaishnav *et al*., 2022) that included 30 and 20 million promoter-driven expression measurements in the yeast *S. cerevisiae* cultured in two media, YPD (complex medium containing yeast extract, peptone, and dextrose) and SD-Ura (synthetic defined medium lacking uracil), respectively. In our study, we neither investigated nor interpreted biological differences between the respective datasets but considered them independent experiments to test the model on additional data.

The train and test datasets were taken as is from the Vaishnav et al. study (Vaishnav *et al*., 2022). A total of 20,616,659 (defined medium) and 30,722,376 (complex medium) random promoter sequences were used to train LegNet in each case. The test data were collected by Vaishnav et al. in independent experiments and included only the high-quality measurements obtained for native (i.e., present in the yeast genome) promoter sequences (3928 for the complex medium, 3977 for the defined medium), see the details in (Vaishnav *et al*., 2022). A subset of the test data from the complex medium was used to compare the performance of LegNet against conventional deep learning methods (DeepSEA, DeepAtt, DanQ). To this end, we used 3733 promoter sequences for which the predictions of DeepSEA, DeepAtt, DanQ, and Vaishnav et al. attention-based model were available in the GitHub repository that accompanied the original yeast GPRA study (Vaishnav *et al*., 2022): https://github.com/1edv/evolution.

To evaluate how LegNet captures the effects of minor sequence alterations, we used the 'genetic drift' data of (Vaishnav *et al*., 2022) where 1 to 3 single-nucleotide substitutions were introduced into 1,000 random starting sequences assessed in both defined and complex media. The respective GPRA data are available in GEO (https://www.ncbi.nlm.nih.gov/geo/) under accession numbers GSE104878 and GSE163045.

### Sequence-to-expression as a soft-classification problem

A straightforward application of machine learning to GPRA experimental data is a regression of a single real value, the expression defined by the cell sorting bin, from a fixed-length DNA sequence. However, such a direct approach cannot fully benefit from domain-specific knowledge and specifics of the GPRA experiment.

We have reformulated the basic sequence-to-expression regression problem as a soft classification task by transforming expression estimates into class probabilities. Given a measured expression *e* (the average of the observed bin numbers), we heuristically assume that the real expression is a normally distributed random variable, see Figure 2b in (de Boer *et al*., 2020):

**Figure 2.**
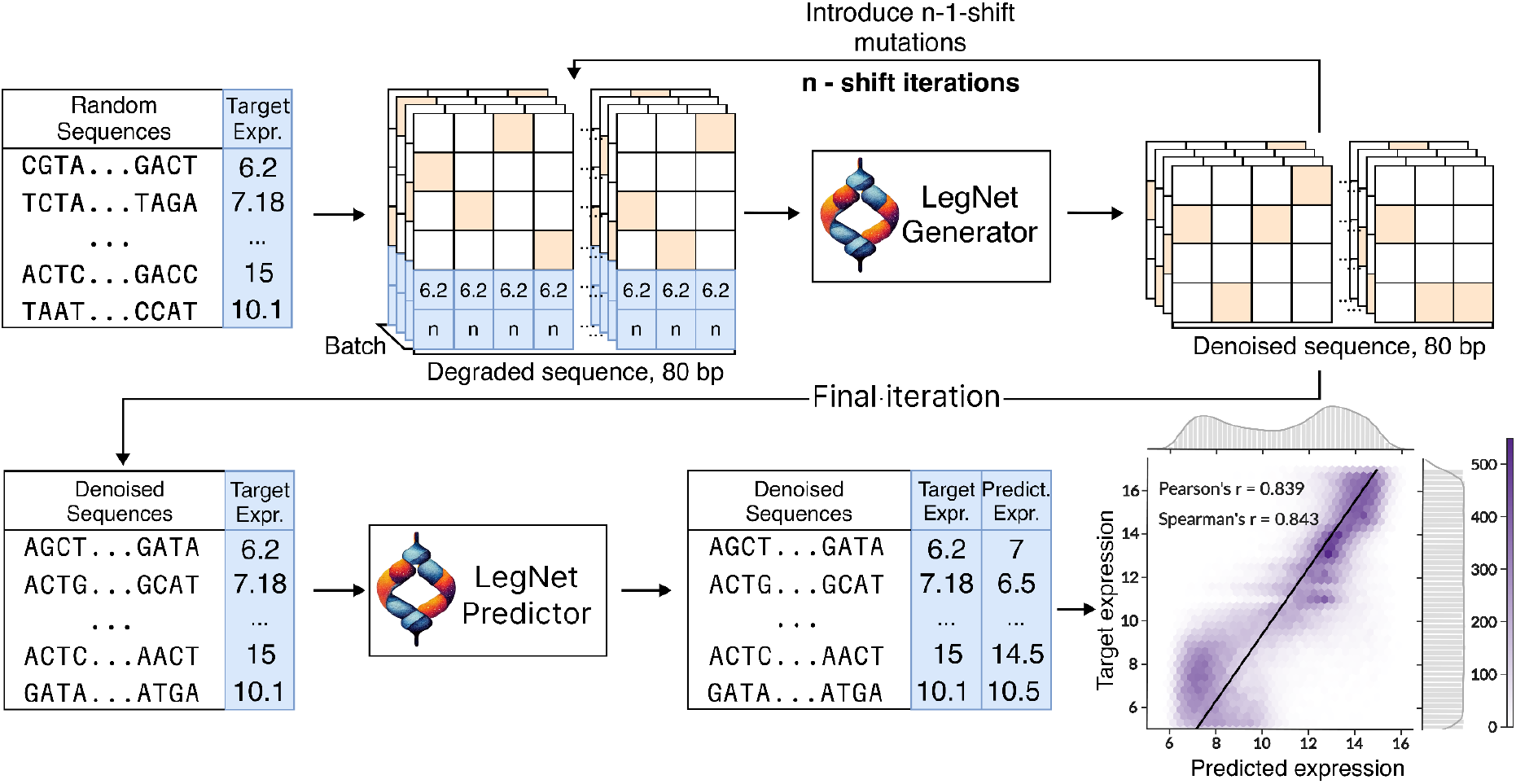
A schematic representation of LegNet application to the rational design of promoters in a flold diffusion framework. The hexagonal binning plot shows the correlation between desired (target) and observed (LegNet-predicted) expression for 110592 designed promoters; the color scale denotes the number of promoters in a bin.

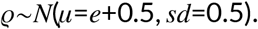

In this approach, for each class *i* from 1 to 16 defined by an original measurement bin, a probability of the class is the cumulative probability to fall into [*i*,*i*+1) range, with 0 and 17 classes (bins) represented by special ranges of (*−*∞, 1] and [17, +∞), respectively.

Thus, for the model loss, we selected the Kullback–Leibler divergence between the distribution derived from the training data and the model output vector containing 18 probabilities corresponding to each class (bin). To obtain a predicted expression value for a sequence during the expression inference step during model validation or test, the predicted probabilities were multiplied by the corresponding bin numbers. This model layer, if joined with softmax, is called soft-argmax (Luvizon *et al*., 2017), see **Figure S1**:

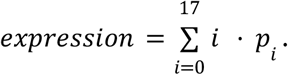

### LegNet architecture

The original LegNet model that won the DREAM2022 challenge (**Figure S1, A**) is based upon a fully-convolutional neural network architecture inspired by EfficientNetV2 (Tan and Le, 2021) with selected features from DenseNet (Huang *et al*., 2018) and additional custom blocks.

The first LegNet block (Stem block) is a standard convolution with `kernel_size=7`, followed by BatchNorm and SiLU activation (**Figure S1, B**). The output of the first block is passed to the sequence of six convolution blocks of EfficientNet-like structure (**Figure S1, C**) but using the grouped convolution instead of the depthwise of the original EfficientNetV2. The standard residual connections are replaced with residual channel-wise concatenation (**Figure S1, C**). Padding mode `'same'` was set for all convolutions. The resize block is of the same structure as the stem block at the start of the network (**Figure S1, B**).

The Squeeze and Excitation (SE) block used as a part of EfficientNet-like block is a modification of that of the original EfficientNetV2 (**Figure S1, E**). The number of parameters in the bilinear block inside of SE block is reduced with low-rank representation of the parameterized tensor via canonical polyadic decomposition implemented in TensorLy (Kossaifi *et al*., 2018) library.

The final block consists of a single point-wise convolutional layer followed by channel-wise Global Average Pooling and SoftMax activation (**Figure S1, D**). We used 256 channels for the first block and [128, 128, 64, 64, 64, 64] channels for six EfficientNetV2-like blocks, respectively. The total number of parameters in the original LegNet model is 1,852,846.

### Adapting and augmenting GPRA data for deep learning

To prepare the data, first, we padded the promoter sequences from the 5' end with the respective constant segments of the plasmids to achieve the total fixed length of 150 bps. Next, sequences were encoded into four-dimensional vectors with one-hot encoding.

We considered the integer expression estimates to belong to the *singleton* promoters observed only once across all bins. The singletons are more likely to have noisier expression estimates, compared to other promoters with non-integer expression values obtained by averaging two or more observations. To supply this information to the model explicitly, we used a binary `is_singleton` channel (1 for singletons, 0 for other training sequences). The final prediction for evaluation used `is_singleton=0`. Since the regulatory elements could be asymmetric with regard to their strand orientation and position relative to the transcription start sites, different scores are expected for the direct and reverse complementary orientation of a particular sequence. Therefore, the training data were augmented by providing each sequence both in native and reverse complementary form, explicitly specifying 0 and 1, respectively, in an additional `is_reverse` channel. We also performed the test-time augmentation by averaging the predictions made for direct (`is_reverse=0`) and reverse complementary (`is_reverse=1`) input of each promoter. A scheme of the input sequence representation is shown in **Figure S2**.

### LegNet training procedure

To train the original model, we used One Cycle Learning Rate Policy (OneCycleLR) (Smith and Topin, 2018) with FastAI (fast.ai - fast.ai—Making neural nets uncool again) modifications: (1) two phases (instead of the original three), (2) the cosine annealing strategy instead of the linear one, (3) the AdamW optimizer (`weight_decay=0.01`) instead of the SGD with momentum. The parameters of the One Cycle Policy were selected using 1/10 of the training data of the DREAM challenge. To select the max learning rate (0.005) for the One Cycle Learning Rate Policy, we used the LR-range test as suggested in (Smith and Topin, 2018).

Each epoch consisted of 1000 batches of size 1024. The model was trained for 150 epochs (defined medium), 300 epochs (complex medium), and 80 epochs (DREAM2022 chardonnay grape juice medium) achieving a reasonable trade-off between training time and validation variance. For the original LegNet model, we used the hyperparameters based on the validation on the last k-fold (10th) of the training data, and the final model was trained from scratch on the whole training dataset.

We used the same weight initialization as in EfficientNetV2 (Tan and Le, 2021). The training of the final model using the NVIDIA RTX A5000 GPU and PyTorch version 1.11.0+cu113 took about 12 hours for the defined medium data, 24 hours for the complex medium data, and 4 hours for the chardonnay grape juice medium data.

### LegNet ablation study and the optimized LegNet model

To identify the key elements of the LegNet architecture, we performed a hierarchical step-by-step analysis using the DREAM2022 train and test GPRA data. We sequentially checked the impact of (1) particular blocks and layers of the network, (2) the choice of an optimization algorithm, (3) the soft-classification approach and the singleton channel, (4) the reverse-complementary augmentation, see the details below. In each case, we trained 5 models with 5 different fixed starting seeds to estimate variability.

We checked the following features of the network architecture for their impact on model performance:

1) Usage of SiLU activation before average pooling;

2) Type of SE-block (comparing to the original EfficientNetV2 block);

3) Type of residual connections (ResidualConcat vs. ResNet Residual block);

4) Different numbers of groups in grouped convolutions;

5) EfficientNetV2 method to estimate the internal dimensionality of the respective block;

6) Overall number of blocks in the model.

The complete list of tested model variants is provided in Supplementary Table 1.

### Optimization algorithm and learning rate schedule

We checked if the AdamW optimizer can be replaced with a recent Lion optimizer (Chen *et al*., 2023), for which the learning rate was tenfold reduced, and the weight decay was tenfold increased, as suggested by the authors. Next, we tested if learning the OneCycleLR scheduler is indeed optimal compared to the widely used ReduceLROnPlateau. As the latter requires the validation subset not only for tuning hyperparameters but during the whole model training, in this comparison we used 10 models trained with 10-fold cross-validation, with the final performance measured on the independent test set.

#### Diffusion model setup for rational promoter design

To adapt LegNet for the rational design of promoter sequences, we used an approach based on cold diffusion (Bansal *et al*., 2022). First, we used the original LegNet on several subsets of GPRA data and, with numeric simulation, estimated the expected number of mutations (300) that is sufficient to make the distribution of expression values indistinguishable from that of random sequences (**Figure S3**). Of note, the mutation process here and below does not have a memory i.e. it does not check if it accidentally reverts back any previously introduced substitutions.

Next, we trained LegNet-Generator (Figure S4) to iteratively correct multiple single-nucleotide substitutions in sequences with known expression values. As input, we supplied a promoter sequence with 0 to 300 mutations, the respective number of mutations, and the expression of the initial sequence. A cross-entropy loss between the reconstructed and original sequence was used during the network training with the following parameters: 200 epochs, AdamW optimizer, learning rate 0.001, batch size 1024, 1000 batches per epoch, and 4 to 1 train-to-validation ratio.

The generation of sequences was performed by the iterative correction of mutations in a cold diffusion manner (**Figure 2**). As input to LegNet-Generator, we supply a random sequence, the number of mutations (0 to 300), and the target expression value. The generated sequence is then changed by introducing `n-1-shift` substitutions and re-supplied to the LegNet-Generator with `n-1` specified as the number of mutations. This cycle is repeated iteratively until the number of introduced mutations does not reach 0. The usage of a non-zero `shift` is necessary to trick the model into introducing more changes and drifting farther away from the starting sequence. Finally, we launched the LegNet-Predictor on the obtained sequences and compared the values against the originally defined expression values. As an illustrative example in this study, we used 100 iterations (with 100 introduced mutations at the first step) and the `shift` of 30, although further optimization of these parameters remains possible.

## Results

### LegNet accurately predicts promoter expression

In this study we present LegNet, a new fully-convolutional neural network architecture inspired by EfficientNetV2 (Tan and Le, 2021), see **Figure S1**. First, we evaluated LegNet in predicting native promoter expression for GRPA data from yeast grown in complex (YPD) or defined (SD-Ura) media. In both cases, LegNet demonstrated consistently high performance, scoring significantly higher than the state-of-the-art transformer model published along with the GPRA data by Vaishnav et al. (Vaishnav *et al*., 2022) (**Figure 3**). Note that the prediction “wall” encountered at around expression levels 4 (complex) and 2.5 (defined) is a known issue with the training data also learned by models of (de Boer *et al*., 2020; Vaishnav *et al*., 2022), which is likely caused by the cell sorter having limited signal-to-noise ratio in this range or inadvertently truncated distribution. We also compared LegNet against earlier deep learning approaches tested in (Vaishnav *et al*., 2022) (**Figure S5**) highlighting the gap between LegNet (~0.96-0.98 Pearson and Spearman correlation against the ground truth test data) and conventional deep learning models such as DeepSEA and DanQ (correlations around ~0.92-0.94).

**Figure 3.**
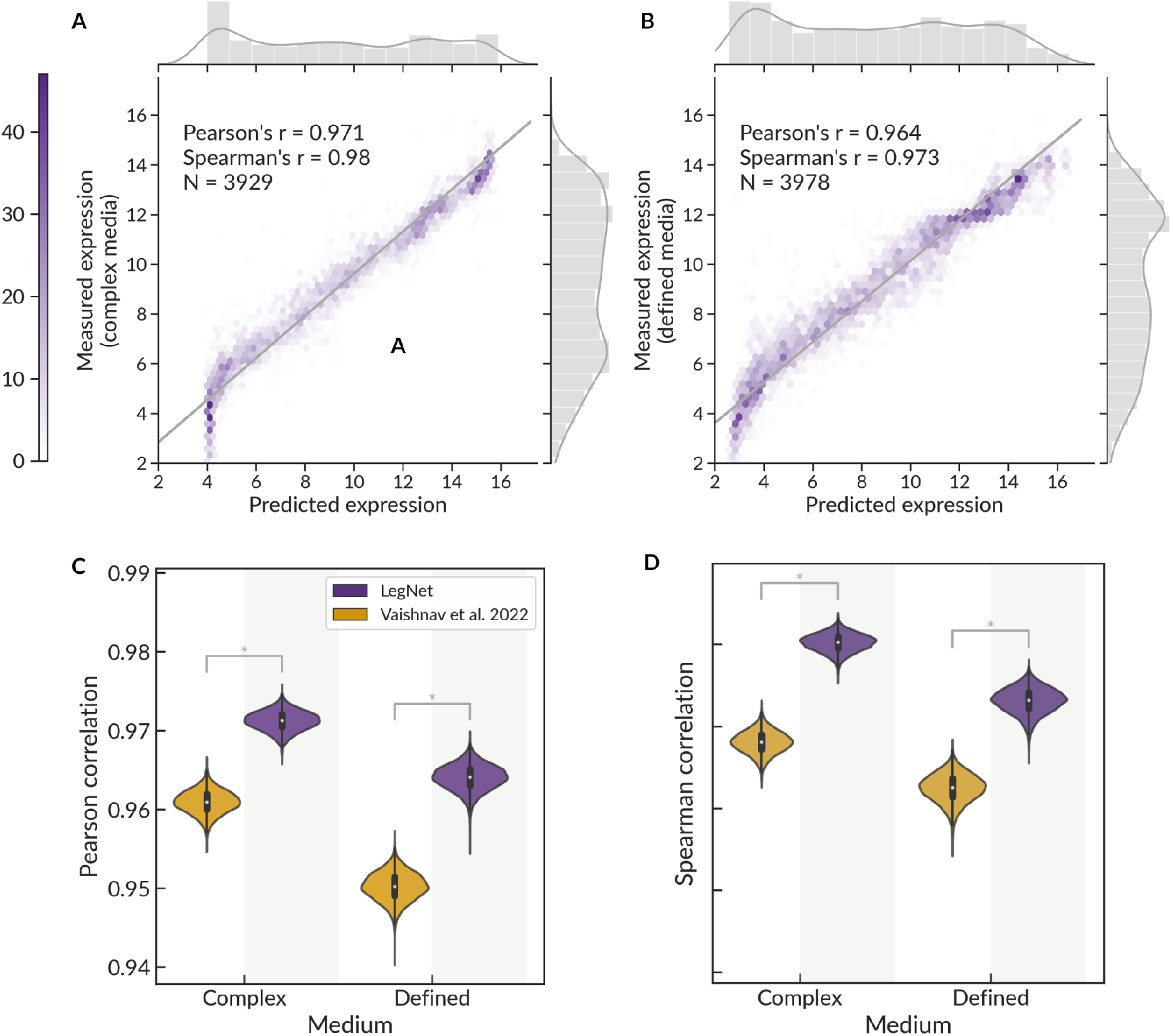
LegNet accurately predicts promoter expression. **A-B**. Prediction of native promoter expression for yeast grown in complex medium (YPD, A) and defined medium (SD-Ura, B), hexagonal binning plots, the color scale denotes the number of promoters in a bin. s**C-D**. Comparison of LegNet prediction performance for native yeast promoter sequences compared to the transformer model of Vaishnav et al.; C: Pearson correlation between predictions and ground truth; D: Spearman correlation; note the Y-axis lower limit. Violin plots show bootstrap with n=10,000. *p < 0.001, Silver dependent correlations test (Silver *et al*., 2004) for the total data.

### LegNet delivers accurate estimates of sequence variant effects

In the DREAM challenge, LegNet was highly successful in estimating the expression of promoters with single-nucleotide variants. To demonstrate it with independent data and further explore LegNet reliability in predicting the effects of multiple nucleotide substitutions, we utilized the GRPA data capturing expression divergence under random genetic drift. For 1,000 unique random promoter sequences, Vaishnav et al. randomly introduced single-nucleotide mutations for three generations and measured the promoter expression in each.

We evaluated the capability of LegNet to quantitatively estimate the difference between expression for original and mutated promoter sequences depending on the number of nucleotide substitutions (1,2, or 3) and compared the performance with the state-of-the-art transformer model of Vaishnav et al. Estimating the single-nucleotide variant effects was the most difficult, but LegNet showed a consistent and significant increase in prediction performance in all scenarios (**Figure 4**).

**Figure 4.**
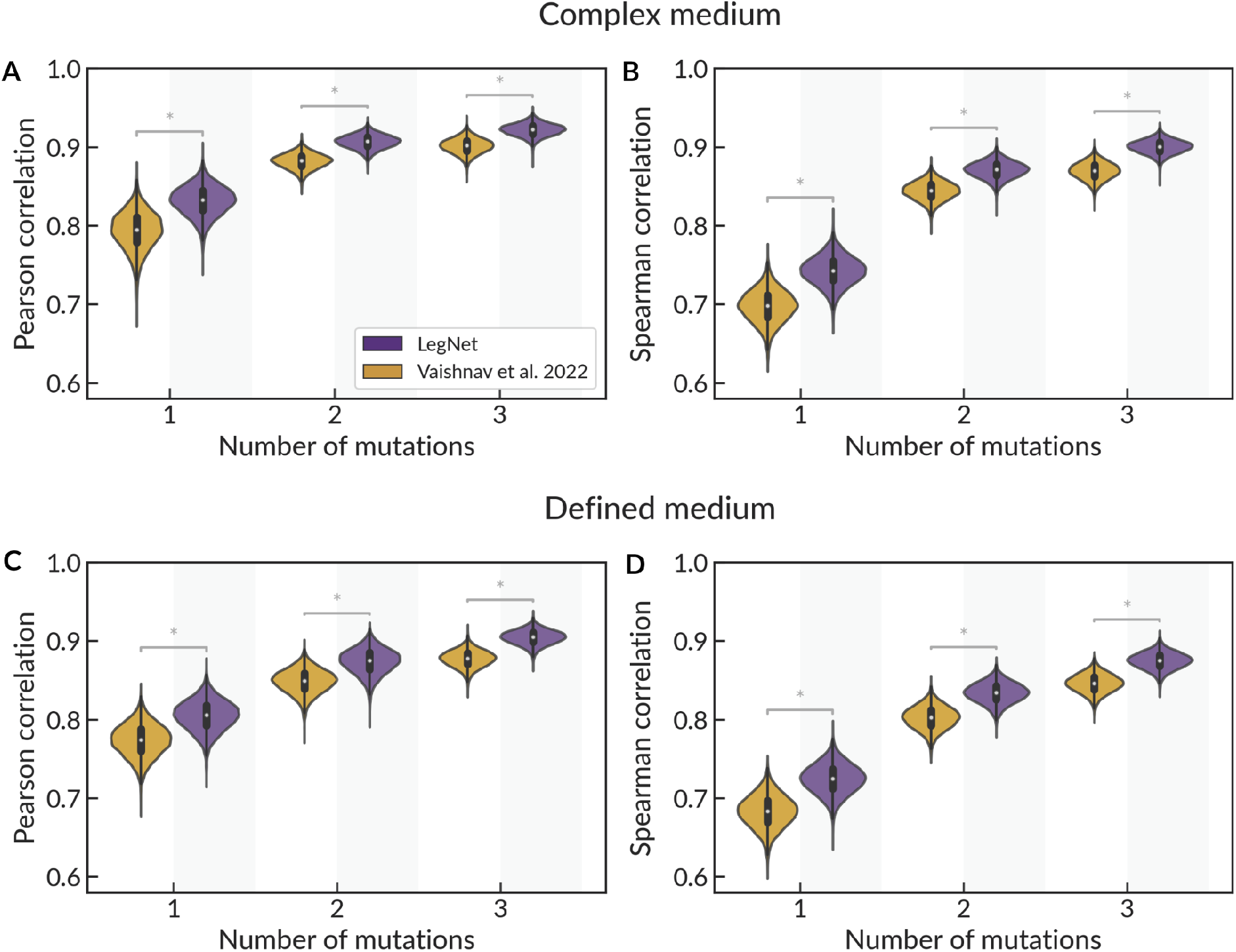
LegNet demonstrates better prediction of variant effects for yeast grown in complex (**A-B**) and defined (**C-D**) medium compared to the transformer model of Vaishnav et al. A, C: Pearson correlation between predictions and ground truth; B, D: Spearman correlation; note the Y-axis lower limit. Violin plots show bootstrap with n=10,000, *p < 0.0001, Silver dependent correlations test (Silver *et al*., 2004) for the total data.

Identifying key performance-affecting elements of LegNet and designing the optimized architecture

By testing different versions of our model (see Methods and Supplementary Table 1), we have found a way to further improve the original LegNet (**Figure S6**) on top of the original DREAM2022 version. The primary architectural changes did not introduce any major changes in the model performance, except for the SE block, which had a major added value of more than +0.005 for both Pearson and Spearman correlations as we optimize around 0.975. The other changes were less impactful and we decided to bring the optimized version of LegNet closer to the original EfficientNetV2.

Thus, the optimized LegNet uses a standard EfficientNetV2 SE-block instead of an originally used custom variant, depthwise instead of grouped convolutions, the original method of EfficientNetV2 to set the dimensionality of the EfficientNetV2-like block, and does not include activation at the final layer before average pooling. However, we kept ResidualConcat as residual connections were important for reaching optimal scores (see Supplementary Table 1). We also kept the original total number of blocks which provide a 79 base pairs receptive field that is close to the actual variable length of the tested promoter. The resulting version of LegNet (2.1M parameters) was used in the consequent tests to prove the importance of soft classification instead of the direct regression of the promoter activity, the singleton channel, and the reverse-complement data augmentation, see **Supplementary Table 1**.

In terms of the optimizer used for the model training, a recently introduced Lion (Chen *et al*., 2023) provided improved performance over AdamW, and in terms of the learning rate scheduler, OneCycleLR was clearly superior compared to the widely used ReduceLROnPlateau. Notably, in terms of the performance differences, the training mode was more important than the minor features of the model architecture, except for the ablation of the SE-block.

### Predictive performance can be further improved on top of the optimized LegNet

On top of the optimized model, we constructed and tested a series of ensemble models by averaging the predictions of 2 to 50 optimized LegNets trained with different starting seeds. Interestingly, the performance of the ensemble stably and significantly increased saturating at 0.9766 and 0.98 (Pearson and Spearman correlations, respectively) for averaged predictions of 50 models compared to 0.9756 and 0.979 of the single optimized LegNet. This suggests that, despite being the best-in-class, LegNet does not reach the theoretical bar set by the level of experimental variability and, with the currently available data, there remain further possibilities for improvements.

### Setting the ground for the large-scale rational design of regulatory sequences

As LegNet provides a best-in-class solution for predictive promoter expression, we further explored its capabilities in the rational design of sequences with a desired level of expression using the diffusion approach. First, we train the LegNet-Generator model to correct the artificial noise by reverting back point mutations introduced in sequences with known expression levels. Next, we perform iterative generation by applying LegNet-Generator to induce substitutions in a completely random sequence, i.e. by tricking the model to correct “errors” in a provided random sequence so that upon full correction the resulting sequence provides a desired expression level. Finally, we verify the results with the LegNet-Predictor based on the original LegNet, see **Figure 2, Supplementary Figure S4**, and Methods. The Pearson and Spearman correlation between the target (as requested) and actually generated (as predicted by the original LegNet) expression reach 0.839 and 0.843, respectively, with imperfect results in a low expression range, but a good agreement between the target and obtained expression for medium-to-highly expressed promoters.

## Discussion

In this study, we presented LegNet, a new deep-learning approach for predicting promoter expression from a DNA sequence. LegNet is an EfficientNetV2-based fully convolutional neural network (Tan and Le, 2021) employing domain-specific ideas and improvements to reach accurate expression modeling and prediction from a DNA sequence.

Multiple factors contribute to the overall LegNet performance, from the up-to-date global model architecture to domain knowledge that facilitated the proper data augmentation and allowed to reformulate the basic regression problem in a way to reflect the scheme of the underlying wet-lab experiment. The data preparation was extended over the standard one-hot encoding approach by discerning whether a target sequence was observed in the experiment only once (a singleton) or multiple times, as the singletons constitute more than half of the training data but eventually provide noisier expression estimates. Next, we augmented the data with reverse complementary sequences. Finally, we reformulated the expression prediction as a soft-classification problem: LegNet was trained to predict not the single expression value but a vector of expression bin probabilities and combine the predicted probabilities into a single predicted expression value at the model evaluation stage.

The exact values of LegNet parameters are likely specific to a particular GPRA experimental setup, so it is recommended to tune the parameters for better applicability of LegNet if trained on data from other experiments or other species, e.g. handling different random insert lengths and adapting the soft classification strategy to different cell sorting techniques and/or cell type-specific experimental variability.

Of note, many studies of machine learning applications in biology attribute the most of the credit to model architecture. However, here we are showing the critical impact of the training scheme on the final model performance. Such an effect is acknowledged in computer vision applications (Bello *et al*., 2021). Also, the performance of a single optimized LegNet model instance was further improved by the LegNet ensemble, suggesting that the potential for building better models using existing GPRA data has not yet been exhausted.

Finally, we have presented a new model to generate promoters with desired expression level, which is a convenient and computationally efficient way compared to the straightforward greedy sequence expression optimization with a predictive model (Wang *et al*., 2020; Kotopka and Smolke, 2020). Previously, convolutional networks have been successfully used for guided selection and design of bacterial and yeast promoters (Wang *et al*., 2020; Kotopka and Smolke, 2020). A more advanced approach was taken for yeast 5'- and 3'-UTRs where Wasserstein generative adversarial network was coupled with a predictive convolutional network (Zrimec *et al*., 2022). However, considering diffusion networks, there are few emerging attempts (Avdeyev *et al*., 2023), and, to our knowledge, the LegNet adaptation is the first generative diffusion framework for promoters built upon the massive data from parallel reporter assays.

All in all, by using the data from GPRAs, we have demonstrated LegNet's efficacy in predicting expression *per se*, quantitatively estimating the effects of sequence variants, and rationally designing promoters with desired expression. We have shown that LegNet shows notably better performance than conventional models and the previous state-of-the-art transformer model. Thus, while today the researchers' preference is biased toward complex architectures, we conclude that the fully convolutional networks should be considered as a reliable approach to the computational modeling of short gene regulatory regions and predicting effects of regulatory sequence alterations.

## Acknowledgments

The study was supported by RSF [20-74-10075 to I.V.K.]. AZ was supported by a personal fellowship from the Non-commercial Foundation for Support of Science and Education “INTELLECT”. We thank Google TRC for providing free access to computational resources used in the model development and assessment. The LegNet logo was generated with Midjourney AI and used under the CC BY-NC license.

