## Supplementary Figures for "LegNet: a best-in-class deep learning model for short DNA regulatory regions"

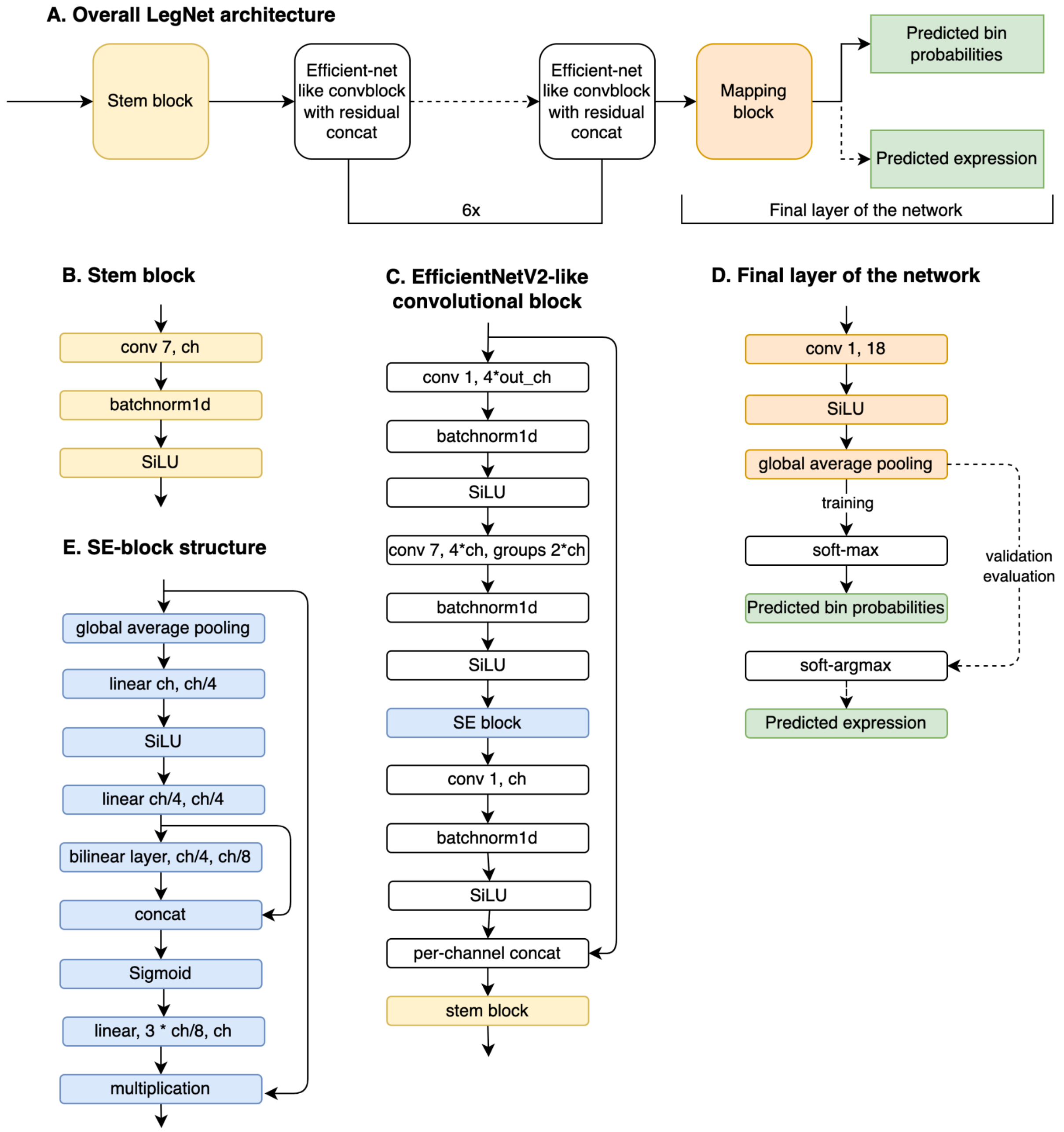


**Figure S1.** LegNet architecture. **A** - overview. **B** - stem block structure. **C** - EfficientNetV2-like convolutional block. **D** - final layer of the network. Dashed lines denote the procedures used during the validation and evaluation only. **E** - SE-block structure. Here `conv k, t*ch` denotes a one-dimensional convolutional layer with kernel size `k` which produces `t * input` channels; `linear t1*ch, t2*ch` denotes a linear layer with `t1` and `t2` coefficients corresponding to the number of input and output channels relative to the input tensor.

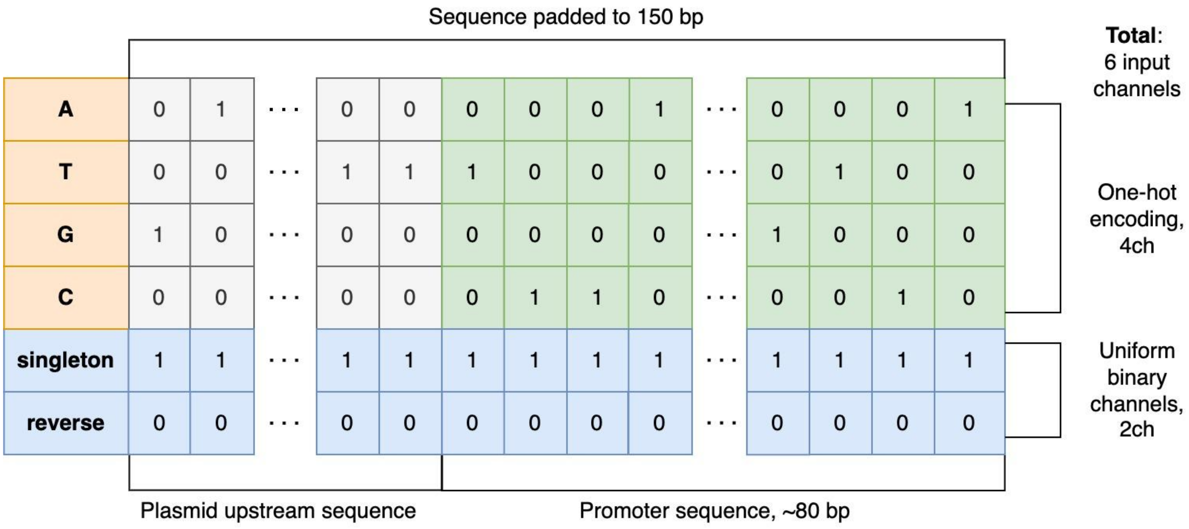


**Figure S2.** LegNet sequence representation. A single sequence is represented position-wise in four channels for one-hot encoded nucleotides, with two extra constant channels signifying whether that supplied sequence was a singleton (i.e. had an integer expression value) and whether the original sequence was subjected to the reverse complementary transformation.

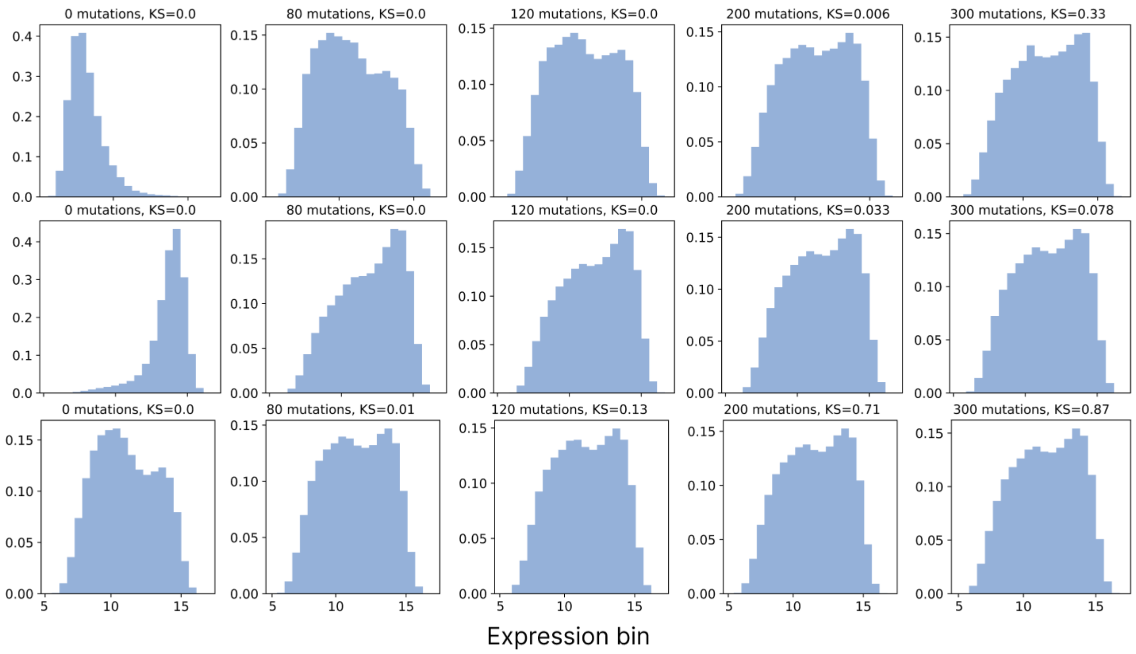


**Figure S3.** Estimating the number of mutations sufficient to change the expression distribution of a starting sequence set into that of a random sequence set. The first column (1): starting expression distributions obtained by randomly sampling 10000 promoters from particular bins of GPRA data. Next columns (2-5): expression distribution after introducing a given number of mutations in each sequence. Top row: sequences sampled from expression bins num. 5-6; middle row: sequences sampled 14-15 expression bins; KS: Kolmogorov-Smirnov test comparing the empirical distribution against predicted expression for a random set of sequences.

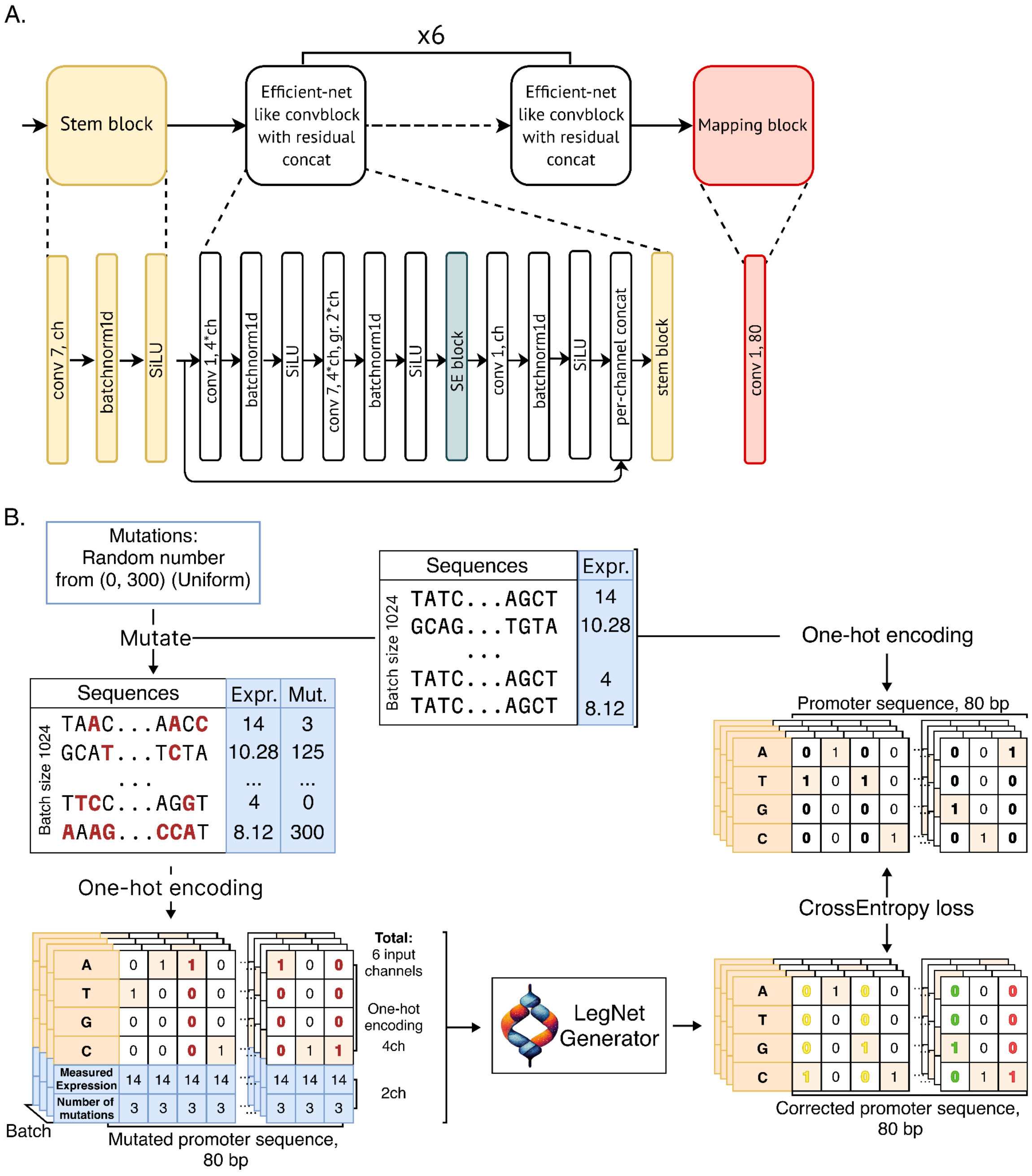


**Figure S4. A.** LegNet-Generator architecture. The principal difference from the original LegNet is the mapping block (highlighted in red). **B.** Training LegNet-Generator to iteratively correct introduced substitutions. In the model output (bottom right) green marks the mutated position that LegNet restored correctly, yellow marks the position unnecessarily changed by LegNet, and red marks the uncorrected mutated position.

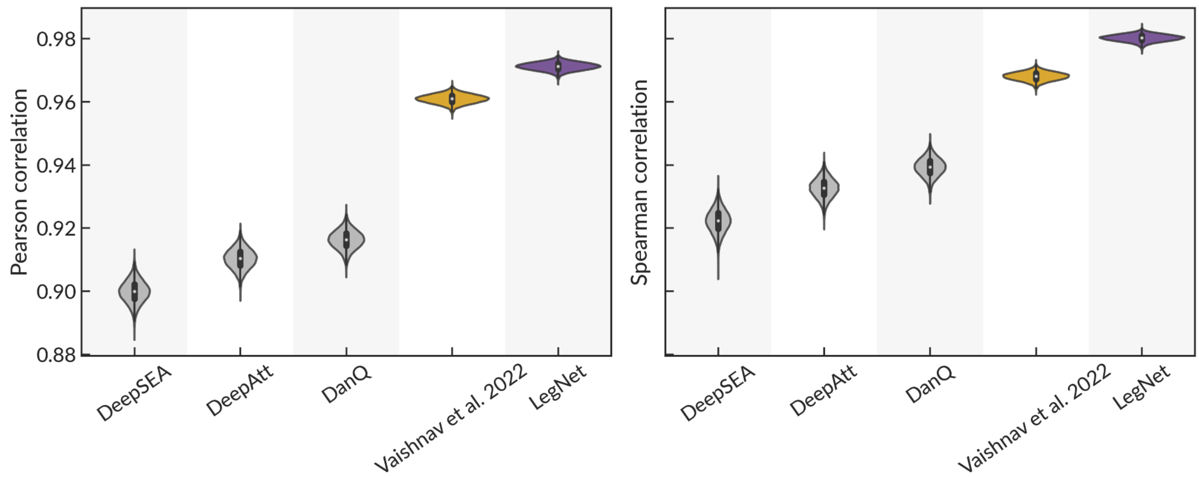


**Figure S5**. LegNet prediction performance for native yeast promoter sequences (complex medium) compared to DeepSEA, DeepAtt, DanQ, and transformer model of Vaishnav *et al.*

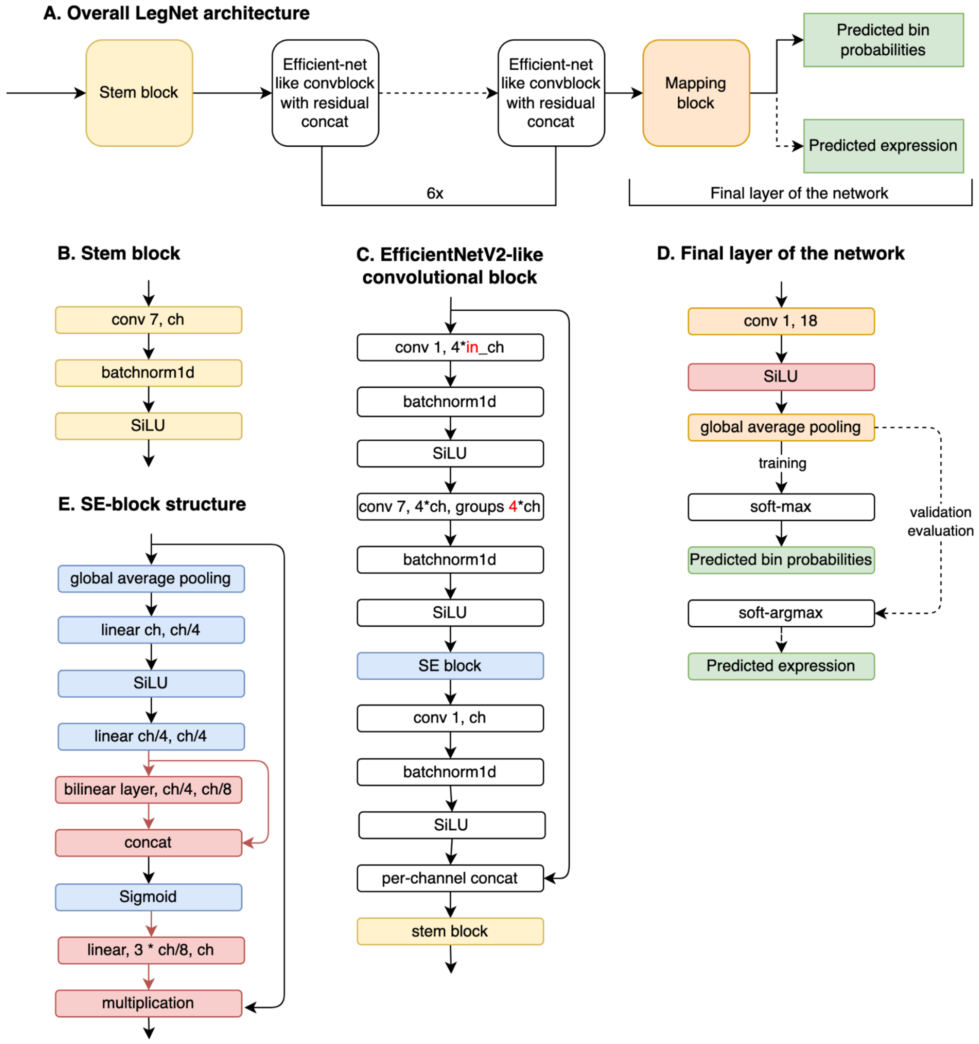


**Figure S6.** Optimized LegNet architecture. Blocks removed from the original architecture are highlighted in red. Red text labels denote modified parameters.

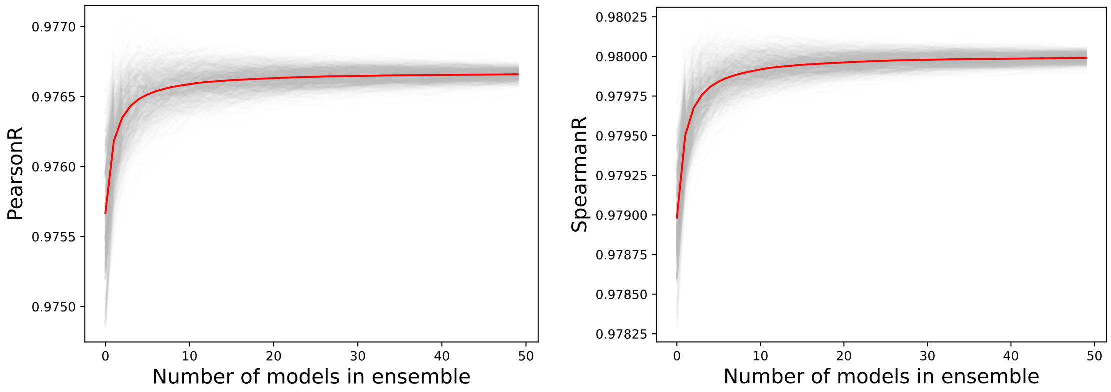


**Figure S7.** Performance of the LegNet ensemble. Left panel: Pearson correlation coefficient, right panel: Spearman correlation coefficient. Gray trajectories: individual samples of a fixed size (X-axis) from a hundred of trained models; red curve: mean value. Note the Y-axis lower limit.

### Supplementary Tables

**Supplementary Table 1.** Detailed results of the LegNet ablation study.
